# LEO1 loss promotes ER stress-adapted migration and cholesterol dependency in colorectal cancer

**DOI:** 10.64898/2026.05.17.725800

**Authors:** Su Chan Park, Ji-Yeon Lee, So Hyun Kwon, Eun Jung Park, Ji Min Lee

## Abstract

The RNA polymerase-associated factor 1 complex (PAF1C) is an evolutionarily conserved transcription elongation complex that regulates RNA polymerase II-mediated transcription and chromatin modification. LEO1, a core subunit of PAF1C, has been implicated in developmental gene regulation, WNT signaling, and leukemogenesis; however, its role in solid tumor progression remains poorly understood. In this study, we found that although LEO1 expression is generally elevated in colorectal cancer (CRC), its expression is reduced in stage IV tumors and is associated with poor clinical outcomes. To investigate its function, we established LEO1 -deficient HCT116 cell line and performed transcriptomic analyses. Loss of LEO1 suppressed epithelial differentiation and developmental gene programs while inducing cell cycle delay. Despite these changes, LEO1-deficient cells exhibited aggressive phenotypes, including enlarged nuclei and increased expression of migration-associated genes, which were further enhanced under glucose deprivation. Motif analysis identified FOXM1 as a key regulator of these migration-related genes. Mechanistically, LEO1 deficiency promoted accelerated transcriptional activation of GRP78, a central regulator of endoplasmic reticulum (ER) stress adaptation. GRP78 was required for survival under ER stress conditions, and its inhibition suppressed both migration and migration-associated gene expression. In addition, transcriptomic analyses revealed upregulation of cholesterol metabolism-related genes in LEO1-deficient cells. Consistently, treatment with the HMG-CoA reductase inhibitor atorvastatin selectively impaired their survival, indicating cholesterol metabolic dependency. Collectively, these findings demonstrate that LEO1 loss promotes ER stress-adapted migration and cholesterol metabolic dependency in CRC, suggesting that these pathways may represent therapeutic vulnerabilities in metastatic LEO1-low CRC.

## Introduction

CRC is one of the leading causes of cancer-related mortality worldwide and remains a major clinical challenge due to metastatic progression and therapeutic resistance. Although advances in surgical resection, chemotherapy, and targeted therapies have improved patient outcomes, the prognosis of patients with advanced-stage or metastatic CRC remains poor (1). Metastasis is responsible for the majority of CRC-related deaths and involves a series of coordinated cellular processes, including epithelial–mesenchymal transition (EMT), enhanced migratory capacity, and adaptation to hostile tumor microenvironmental conditions (2, 3). Increasing evidence suggests that metabolic stress within the tumor microenvironment is a critical driver of these adaptive phenotypes (4). Tumor cells are continuously exposed to nutrient deprivation (5), hypoxia (6), oxidative stress (7), and imbalances in glucose (8) and lipid availability (9), all of which impose strong selective pressures during tumor progression (10). To survive under these conditions, cancer cells undergo transcriptional and metabolic reprogramming that promotes stress adaptation, cellular plasticity, and metastatic competence.

The RNA polymerase-associated factor 1 complex (PAF1C) is an evolutionarily conserved transcription elongation complex that regulates RNA polymerase II-mediated transcription and chromatin modification across eukaryotes (11). LEO1, a core subunit of PAF1C, has been implicated in transcriptional regulation associated with histone modifications, chromatin remodeling, and maintenance of cellular homeostasis. Studies in yeast and fibroblast systems have demonstrated that LEO1 contributes to transcriptional and chromatin adaptation under nutrient stress and quiescent conditions (12–14). Notably, loss of LEO1 induces sterol uptake genes and alters chromatin status (15), suggesting a functional link between transcriptional regulation and metabolic adaptation. However, despite these findings, the functional role of LEO1 in solid tumor progression, particularly under metabolically dynamic conditions within the tumor microenvironment, remains poorly understood.

Metabolic stress has emerged as a hallmark of aggressive tumor progression (16–18). In rapidly proliferating tumors such as CRC, insufficient vascularization and high metabolic demand generate regions of hypoxia and nutrient deprivation. Under these conditions, cancer cells activate adaptive transcriptional programs that suppress proliferation-associated pathways while promoting survival, migration, and stress tolerance (19, 20). Metabolic stress also promotes EMT-like phenotypes and invasive behavior, linking stress adaptation to metastasis. These adaptive responses are closely associated with activation of stress-responsive signaling pathways, including the unfolded protein response (UPR) and endoplasmic reticulum (ER) stress signaling (21). Among ER stress regulators, glucose-regulated protein 78 (GRP78/HSPA5) functions as a master regulator of adaptive UPR signaling and is frequently upregulated in cancer cells exposed to metabolic stress (22). GRP78 supports proteostasis, stress tolerance, metastasis, and therapeutic resistance in multiple cancers (23).

Increasing evidence indicates that metabolic rewiring in cancer extends beyond glucose metabolism and involves profound alterations in lipid and cholesterol metabolism (24). Cholesterol supports membrane integrity, intracellular signaling, and oxidative stress buffering (25). Dysregulation of cholesterol metabolism has been increasingly recognized as a feature of aggressive tumors, where enhanced cholesterol uptake, synthesis, and turnover support survival and metastatic adaptation (26–28). Because ER homeostasis and cholesterol metabolism are tightly interconnected (21), cancer cells exploit these pathways to survive under metabolic stress. These suggest that co-targeting stress adaptation and cholesterol dependency may provide a therapeutic strategy.

In this study, we investigated the role of LEO1 in colorectal cancer progression under metabolic stress conditions. We demonstrate that LEO1 expression is dynamically regulated during CRC progression and is reduced in metastatic CRC and associated with poor clinical outcomes. Functional analyses revealed that loss of LEO1 promotes migratory and EMT-like phenotypes while simultaneously enhancing ER stress adaptation through GRP78 activation. Furthermore, LEO1-deficient CRC cells exhibited metabolic dependency on cholesterol metabolism and increased vulnerability to combined inhibition of cholesterol biosynthesis and ER stress adaptation. Collectively, our findings identify LEO1 as a critical regulator linking transcriptional regulation, stress adaptation, and metabolic rewiring in colorectal cancer and suggest a potential therapeutic strategy for metastatic LEO1-low CRC.

## Results

### LEO1 exhibits context-dependent expression during colorectal cancer progression

Most current insights into LEO1 function have been derived from model organisms or non-transformed cell systems, particularly in the context of cellular quiescence, and its role in solid tumors remains largely unexplored. To investigate the clinical and biological relevance of LEO1 in CRC, we systematically analyzed its expression across patient tissues, CRC cell lines, and publicly available transcriptomic datasets.

Analysis of CRC patient tissues revealed heterogeneous but frequently elevated LEO1 expression in tumor tissues compared with matched normal tissues (**Figure 1A**). Consistent with these observations, most CRC cell lines exhibited increased LEO1 expression relative to the normal colon epithelial cell line HCEC-1CT (**Figure 1B**), suggesting that LEO1 upregulation is a common feature of colorectal tumor cells. To further validate these findings in a larger patient cohort, we analyzed data from the TCGA-COAD cohort. Consistent with the observations in patient tissues and CRC cell lines, LEO1 expression was significantly higher in tumors than in normal tissues (**Figure 1C**). These findings are consistent with previous reports showing that LEO1 amplification is particularly elevated in colorectal cancer compared with other cancer types (29).

**Figure 1.**
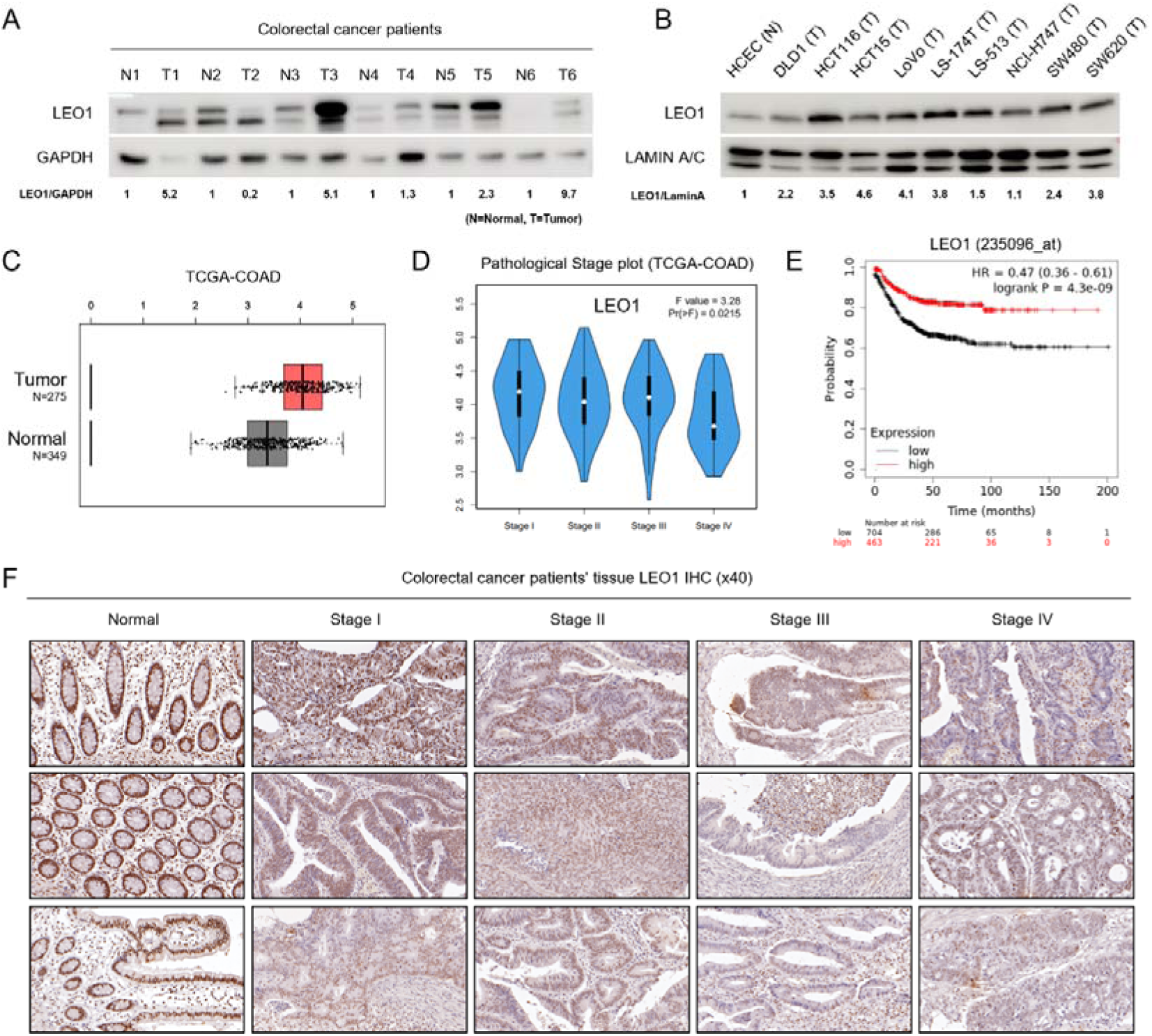
Context-dependent expression and clinical relevance of LEO1 in colorectal cancer. **(A)** LEO1 protein expression was assessed by Western blot analysis comparing paired normal and tumor tissues from CRC patients. **(B)** LEO1 protein expression in a panel of colorectal cancer cell lines was analyzed by Western blot and compared with a normal colon epithelial cell line (HCEC-1CT). **(C)** Analysis of LEO1 mRNA expression in colorectal cancer was performed using the GEPIA2 platform based on the TCGA-COAD cohort. **(D)** LEO1 mRNA expression was stratified according to pathological stage using the GEPIA2 platform based on the TCGA-COAD cohort. **(E)** Kaplan–Meier survival analysis of colorectal cancer patients based on LEO1 expression levels was performed using the Kaplan–Meier Plotter database. **(F)** Representative images of LEO1 immunohistochemical (IHC) staining in formalin-fixed paraffin-embedded (FFPE) colorectal tissues, including normal colon and tumor samples from Stage I to Stage IV patients (n = 3 per stage). LEO1 staining was visualized using DAB (brown), and nuclei were counterstained with hematoxylin (blue). All images were obtained at 40× magnification using CaseViewer.

However, stratification by pathological stage revealed a progressive decrease in LEO1 expression with advancing tumor stage (**Figure 1D**). This stage-dependent reduction suggests that although LEO1 may be initially upregulated during tumor initiation or early progression, its expression becomes diminished as tumors acquire more aggressive phenotypes. Additionally, Kaplan–Meier survival analysis demonstrated that lower LEO1 expression was significantly associated with poorer overall survival in CRC patients (**Figure 1E**). This inverse correlation between LEO1 expression and patient prognosis suggests that loss of LEO1 may contribute to disease progression and the acquisition of aggressive tumor characteristics, enhanced invasiveness, metastatic potential, or adaptation to stress conditions within the tumor microenvironment.

To further evaluate the spatial distribution of LEO1 expression in colorectal cancer tissues, we analyzed LEO1 expression in FFPE CRC specimens. Strong LEO1 expression was observed in the epithelial compartment of normal colonic tissues, whereas gradual decrease in LEO1 expression in LEO1 expression or increases in the proportion of LEO1-low cells were observed from early-stage to advanced-stage tumors (**Figure 1F**). The discrepancy between overall tissue-level expression patterns and histological observations may reflect differences in cellular composition and spatial heterogeneity within tumor tissues.

Collectively, these findings suggest that LEO1 expression is dynamically regulated during colorectal cancer progression. Although LEO1 expression is elevated in a substantial subset of CRC tissues and cell lines compared with normal controls, its expression progressively declines in advanced-stage tumors and is associated with poor patient prognosis. These observations suggest that loss of LEO1 may confer adaptive advantages during CRC progression, thereby contributing to the emergence of aggressive and stress-adaptive tumor phenotypes.

### LEO1 depletion promotes metastatic and EMT-like features

We next investigated the clinical relevance of LEO1 using a Stage IV colorectal cancer specimen with matched liver metastatic tissue. Immunohistochemical analysis of the primary tumor revealed marked intratumoral heterogeneity, with distinct populations of tumor cells exhibiting either high or low LEO1 expression. However, liver metastatic lesions were predominantly composed of LEO1-low cancer cells, resembling the LEO1-low subpopulation observed in the primary tumor (**Figure 2A**). These findings suggest that reduced LEO1 expression may provide a selective advantage during metastatic progression and may be associated with enhanced migratory or stress-adaptive properties.

**Figure 2.**
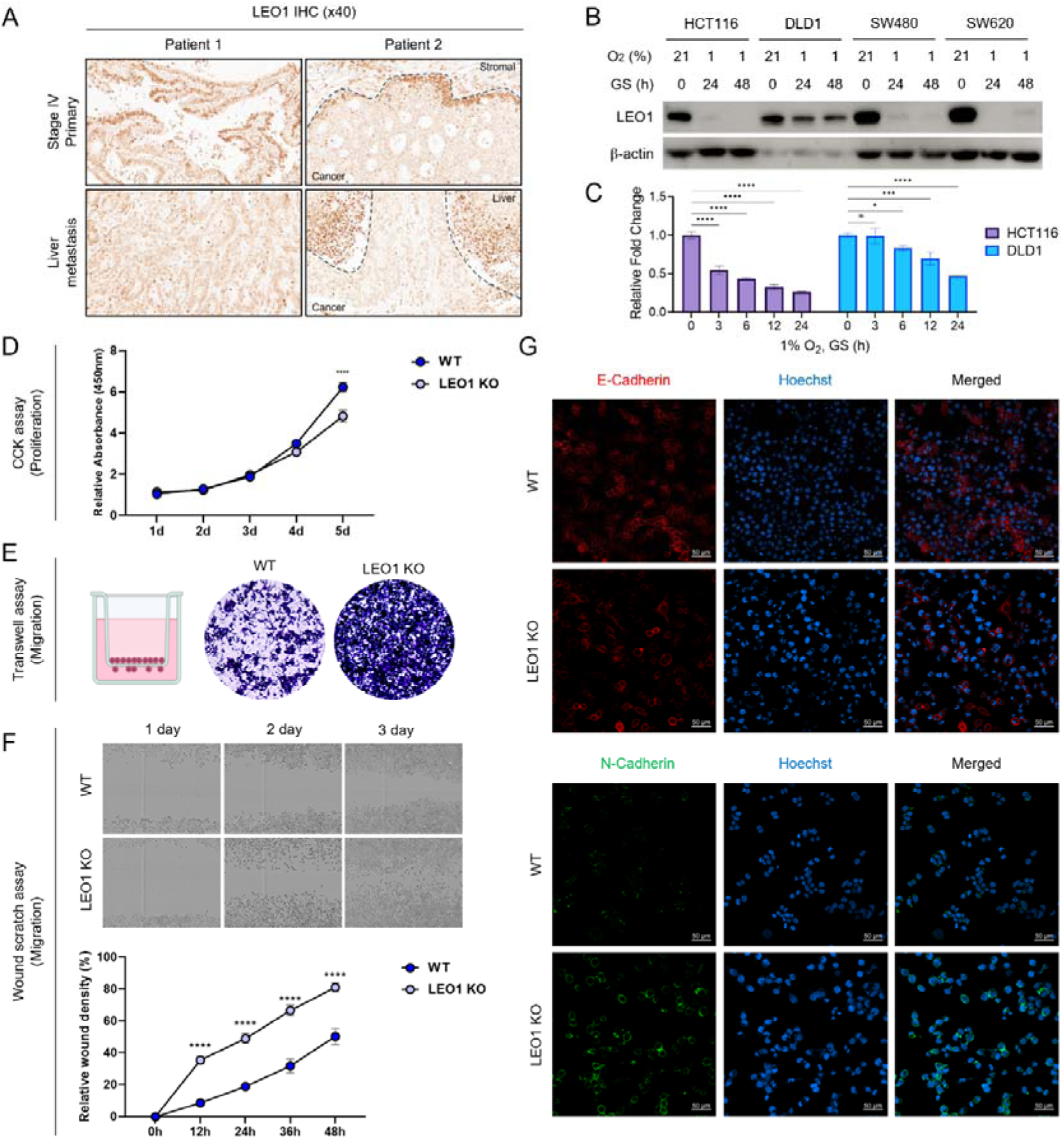
LEO1 loss promotes the migratory capacity of CRC cells. **(A)** Representative IHC staining of LEO1 in colorectal cancer patient tissues, including primary tumors and matched liver metastatic tissues. LEO1 was visualized by DAB staining (brown). All images were acquired at 40× magnification using CaseViewer. **(B)** Western blot analysis of LEO1 protein levels in colorectal cancer cells under TME-mimicking double stress conditions. **(C)** qRT-PCR analysis of LEO1 mRNA levels in colorectal cancer cells under TME-mimicking double stress conditions. Data are presented as mean ± SD. Statistical significance was determined by two-way ANOVA (*p < 0.05, **p < 0.01, ***p < 0.001, ****p < 0.0001). **(D)** Cell proliferation was assessed using a CCK-8 assay in WT and LEO1 KO cells. Data are presented as mean ± SD, and statistical significance was determined by two-way ANOVA (****p < 0.0001). **(E)** Transwell migration assay was performed in WT and LEO1 KO cells. Representative images stained with crystal violet are shown. **(F)** Wound healing assay was performed in WT and LEO1 KO cells. Data are presented as mean ± SD, and statistical significance was determined by two-way ANOVA (*p < 0.05, **p < 0.01, ***p < 0.001, ****p < 0.0001). **(G)** Immunofluorescence staining of epithelial and mesenchymal markers in WT and LEO1 KO cells. Scale bars, 50 μm.

We then sought to model tumor microenvironmental stress conditions *in vitro* by exposing CRC cell lines to combined hypoxia (1% OL) and glucose starvation, thereby inducing metabolic stress. Under these conditions, LEO1 expression decreased in a time-dependent manner at both the protein and mRNA levels (**Figure 2B-C**). This dynamic reduction indicates that LEO1 expression is responsive to environmental stress cues and may be actively suppressed as part of a stress-adaptive transcriptional program. Notably, this reduction was particularly prominent in mesenchymal-like CMS4-type colorectal cancer cell lines, including HCT116, SW480, and SW620.

Building upon our observations that LEO1 expression is downregulated under tumor microenvironmental conditions and associated with metastatic progression, we next investigated the functional consequences of LEO1 loss in CRC cells. To directly evaluate the role of LEO1, HCT116 LEO1 knockout (KO) cells were established, enabling systematic characterization of phenotypic alterations resulting from LEO1 depletion in a controlled cellular context.

Functional analyses revealed that loss of LEO1 induces a phenotypic shift in CRC cells. Cell proliferation analysis demonstrated that LEO1 KO cells exhibited a modest reduction in proliferative capacity compared with wild-type (WT) controls, suggesting that LEO1 partially contributes to basal cell growth (**Figure 2D**). In contrast, migration assays revealed a marked increase in the migratory capacity of LEO1 KO cells, indicating a shift toward a more motile phenotype. Consistently, wound healing assays confirmed that LEO1 KO cells displayed significantly accelerated wound closure compared with WT cells (**Figure 2E-F**).

We next examined whether these phenotypic alterations were associated with epithelial-mesenchymal transition (EMT) related changes (30). Immunostaining analysis demonstrated that LEO1 KO cells exhibited reduced expression of E-cadherin, a key epithelial adhesion molecule, together with increased expression of N-cadherin, a mesenchymal marker associated with enhanced motility and invasiveness (**Figure 2G**). These molecular alterations indicate a shift toward a mesenchymal-like state and suggest that loss of LEO1 promotes EMT-associated transcriptional reprogramming.

Taken together, these findings demonstrate that metabolic stress conditions within the tumor microenvironment may promote LEO1 downregulation in CRC cells, particularly in mesenchymal-like CMS4-type cells. Although LEO1 depletion reduced cell proliferation, it markedly enhanced migratory capacity and induced EMT-like characteristics. Combined with the clinical and histopathological observations, these results suggest that dynamic regulation of LEO1 expression plays an important role in metastatic competence and aggressive tumor behavior in CRC.

### LEO1 loss reduced proliferation but shows nuclear enlargement associated with aggressive features in colon cancers

Given the reduced proliferation and enhanced migratory capacity observed in LEO1 KO cells, we next investigated the underlying molecular mechanisms through transcriptomic analysis. RNA sequencing was performed under basal conditions to compare WT and LEO1 KO cells, to identify intrinsic transcriptional alterations independent of external stress conditions. Gene ontology (GO) enrichment analysis revealed global downregulation of genes associated with epithelial cell proliferation and development in LEO1 KO cells (**Figure 3A**). Consistently, heatmap analysis demonstrated coordinated suppression of genes associated with the GO terms “positive regulation of epithelial cell proliferation” and “epithelial cell development” in LEO1 KO cells (**Figure 3B**). In addition, several growth-associated genes implicated in colorectal cancer progression, including WNT3A and MYC (31), were significantly reduced following LEO1 loss. These findings suggest that LEO1 contributes to the maintenance of transcriptional programs required for epithelial growth and proliferative signaling.

**Figure 3.**
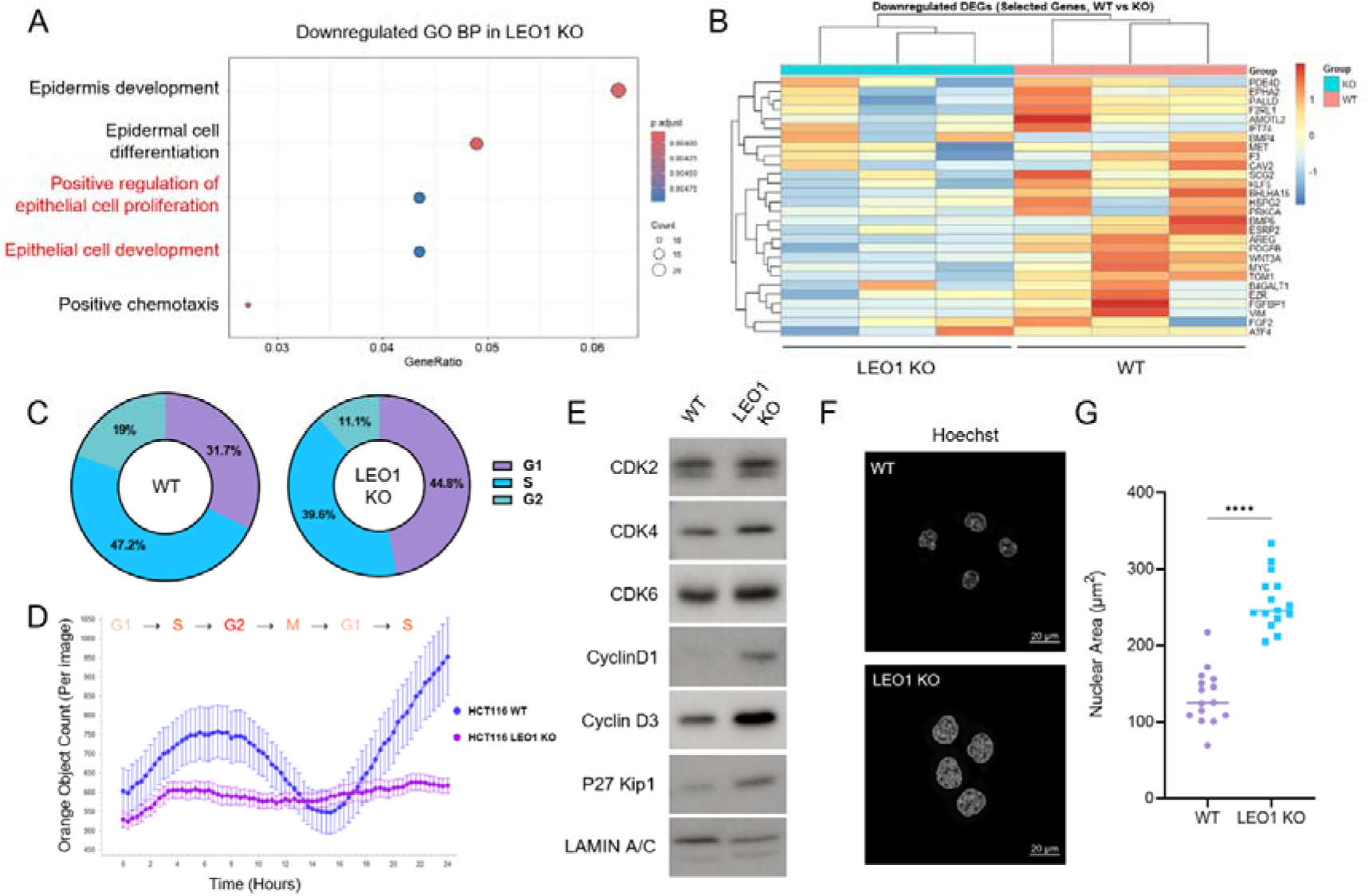
LEO1 KO is associated with reduced proliferation but exhibits an aggressive nuclear enlargement phenotype. **(A)** GSEA GO BP enrichment analysis of downregulated genes in LEO1 KO cells. **(B)** Heatmap showing downregulation of genes associated with the GO terms “positive regulation of epithelial cell proliferation” and “epithelial cell development” in LEO1 KO cells compared with WT cells. **(C)** Cell cycle distribution analysis by propidium iodide staining in HCT116 WT and LEO1 KO cells. **(D)** Live-cell imaging analysis of cell cycle progression using a geminin reporter. **(E)** Western blot analysis of cell cycle-related proteins in WT and LEO1 KO cells. **(F)** Representative Hoechst-stained images showing enlarged nuclei in HCT116 LEO1 KO cells compared with WT controls. Scale bars, 20 μm. **(G)** Quantification of nuclear area in HCT116 WT and LEO1 KO cells. Nuclear area was measured using ZEN 3.6 software. Statistical significance was determined by an unpaired Student’s t-test (****p < 0.0001).

To further investigate the functional consequences of these transcriptional alterations, we analyzed cell cycle progression in LEO1 KO cells. Cell cycle profiling revealed a marked shift in cell cycle distribution, characterized by accumulation of cells in the G1 phase together with a reduction in S and G2 populations compared with WT cells (**Figure 3C**). This distribution pattern is indicative of impaired cell cycle progression and suggests delayed or restricted transition from G1 to S phase. Consistent with this observation, live-cell imaging using a geminin reporter demonstrated delayed progression through S/G2 phases in LEO1 KO cells (32), providing additional evidence of altered cell cycle dynamics following LEO1 depletion (**Figure 3D**). At the molecular level, loss of LEO1 was associated with altered expression of key cell cycle regulators, including CDK2, CDK4, CDK6, Cyclin D1, and Cyclin D3, together with increased expression of the CDK inhibitor p27^Kip1^ (**Figure 3E**). These molecular changes are consistent with a altered cell cycle state and further support the role of LEO1 as a positive regulator of cell cycle progression in colorectal cancer cells.

Interestingly, despite the reduced proliferative capacity and cell cycle transition, LEO1 KO cells exhibited a marked increase in nuclear size compared with WT cells (**Figure 3F**). Quantitative analysis confirmed significant enlargement of nuclear area following LEO1 depletion (**Figure 3G**). In CRC, increased nuclear size is associated with chromatin decompaction, altered nuclear stiffness, enhanced invasiveness, and metastatic potential, and is often correlated with advanced tumor stage (33). Therefore, the enlarged nuclear morphology observed in LEO1 KO cells may reflect structural and epigenetic alterations associated with a more deformable and invasive cellular state.

Taken together, these findings demonstrate that loss of LEO1 impairs transcriptional programs involved in epithelial proliferation and cell cycle progression while simultaneously promoting nuclear enlargement associated with aggressive tumor features. These results suggest that LEO1 depletion uncouples proliferative capacity from structural characteristics linked to tumor plasticity and invasiveness, supporting a context-dependent role for LEO1 in regulating colorectal cancer progression.

### LEO1 loss is associated with activation of migration-related programs through FOXM1-dependent transcriptional regulation

Glucose starvation is a common metabolic stress encountered by tumor cells within the tumor microenvironment due to inadequate vascularization and high metabolic demand (34). Under these conditions, cancer cells undergo transcriptional reprogramming to promote survival, cellular plasticity, and invasive behavior. To investigate how LEO1 deficiency influences stress-adaptive responses, we performed RNA sequencing under glucose starvation. GSEA GO BP analysis revealed significant upregulation of pathways associated with cell migration, motility, and locomotion in LEO1 KO cells compared with WT cells (**Figure 4A**). Consistently, GSEA further demonstrated enrichment of migration-related transcriptional programs in LEO1 KO cells under glucose starvation (**Figure 4B**). Heatmap analysis additionally revealed increased expression of genes associated with cell migration, motility, and locomotion following glucose starvation, suggesting activation of a stress-adaptive migratory transcriptional state in LEO1 KO cells (**Figure 4C**).

**Figure 4.**
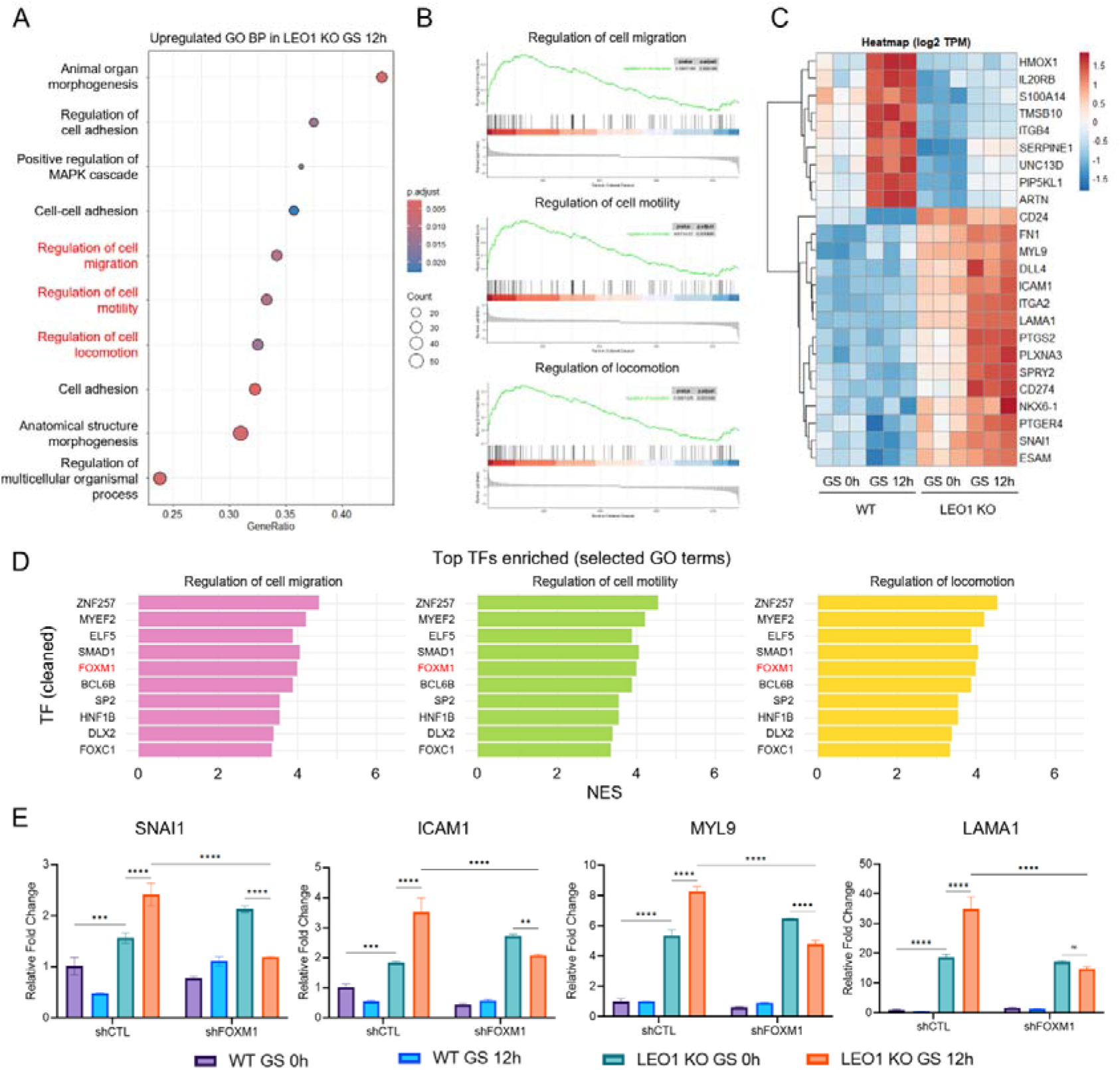
FOXM1 drives migration-related transcriptional programs in LEO1 KO cells under glucose starvation. **(A)** GSEA GO BP analysis showing increased migration-related gene expression in LEO1 KO cells under glucose starvation. **(B)** GSEA plot showing enrichment of migration-related gene sets in LEO1 KO cells. **(C)** Heatmap showing the expression of selected migration-related genes responsive to glucose starvation. **(D)** Bar plot showing transcription factors predicted to regulate migration-related genes as predicted by cisTarget analysis. Enrichment is represented by normalized enrichment score (NES). **(E)** qRT-PCR analysis of representative migration-related genes in WT and LEO1 KO cells following FOXM1 knockdown under glucose starvation conditions.

To identify the transcriptional regulators underlying these changes, motif enrichment analysis using cisTarget was performed on migration-associated gene sets. Among the candidate transcriptional regulators, FOXM1 was consistently identified as a prominent candidate regulator (**Figure 4D**). FOXM1 is a well-established regulator of cancer progression and has been implicated in EMT, invasion, and metastasis (35). Functional validation demonstrated that FOXM1 knockdown significantly attenuated the induction of migration-associated genes under glucose starvation conditions (**Figure 4E**), indicating that FOXM1 is required for activation of the stress-adaptive migratory transcriptional program.

Taken together, these findings demonstrate that LEO1 deficiency promotes FOXM1-dependent transcriptional reprogramming toward a migratory phenotype under metabolic stress conditions. Furthermore, the enhanced dependence on FOXM1 in LEO1 KO cells suggests that loss of LEO1 sensitizes colorectal cancer cells to stress-induced migratory phenotype, thereby contributing to aggressive tumor behavior.

### ER stress response in LEO1 KO cells promotes migration-related transcriptional programs under glucose starvation

While FOXM1-dependent transcriptional reprogramming promoted a migratory phenotype under glucose starvation, the mechanisms by which LEO1 KO cells maintain survival under metabolic stress remained unclear. To address this question, we performed comparative transcriptomic analysis across four experimental conditions: WT and LEO1 KO cells cultured under either high-glucose or glucose starvation conditions. GSEA comparison revealed significant enrichment of ER stress response–related pathways in LEO1 KO cells under glucose starvation (**Figure 5A**). Consistently, heatmap analysis demonstrated increased expression of ER stress–associated genes following glucose starvation in both WT and LEO1 KO cells; however, distinct transcriptional response patterns were observed between the two groups. WT cells preferentially induced ATF4 and DDIT3 (CHOP), which are commonly associated with stress-induced apoptotic signaling (36). In contrast, LEO1 KO cells exhibited stronger induction of adaptive UPR-related genes, including ATF6, EIF2AK3 (PERK), XBP1, and HSPA5 (GRP78) (37–39), suggesting preferential activation of ER stress adaptation programs that support proteostasis and cellular survival (**Figure 5B**).

**Figure 5.**
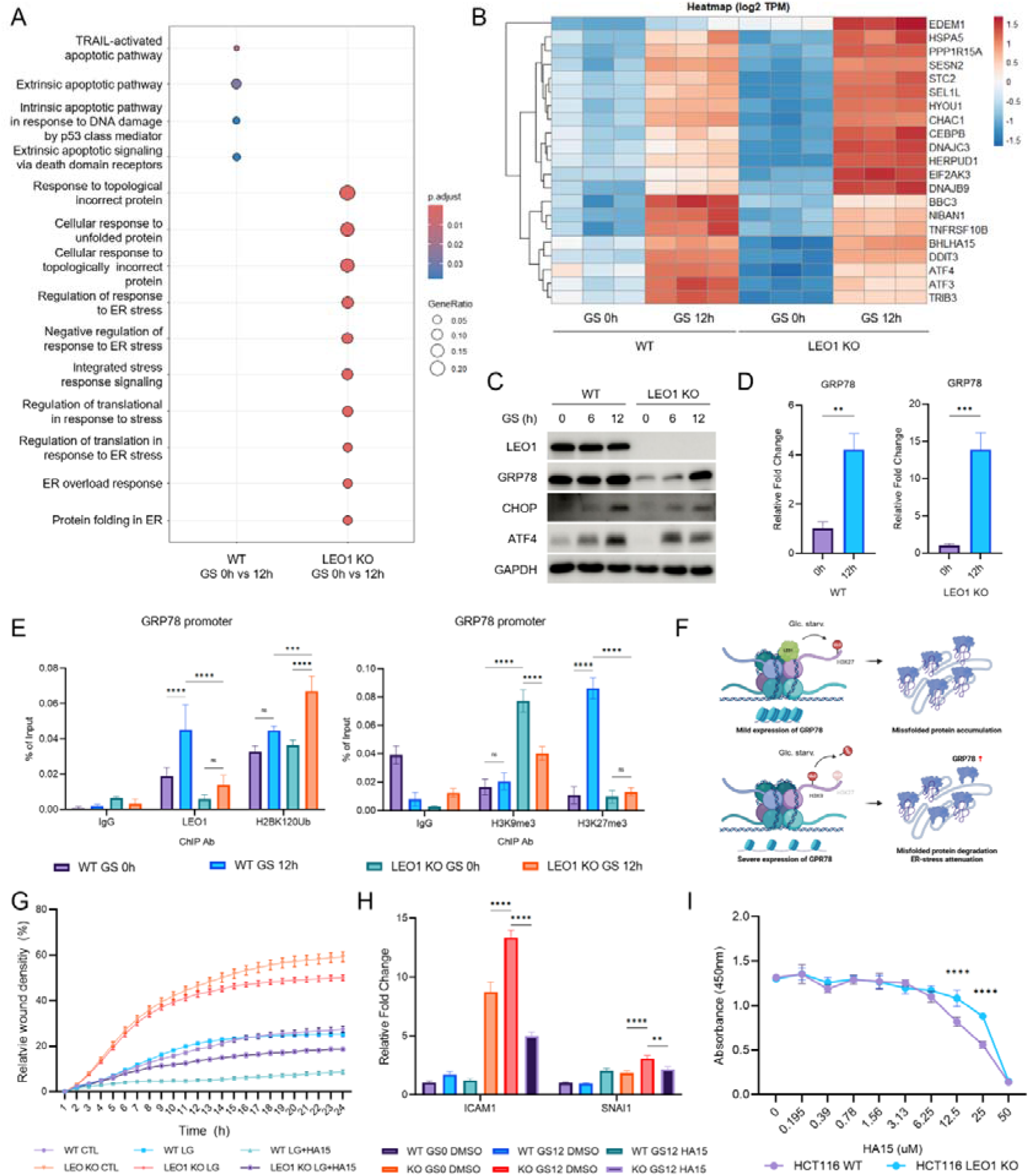
LEO1 deficiency promotes adaptive ER stress responses through epigenetic activation of the GRP78 promoter under glucose starvation. **(A)** GSEA GO BP comparison under glucose starvation conditions in WT and LEO1 KO cells. **(B)** Heatmap showing the expression of genes associated with the GO term “response to ER stress” under glucose starvation. **(C)** Western blot analysis of ER stress-related proteins, including GRP78, ATF4, and CHOP in WT and LEO1 KO cells under glucose starvation. **(D)** qRT-PCR analysis of GRP78 expression in WT and LEO1 KO cells under glucose starvation. Statistical significance was determined by two-way ANOVA (*p < 0.05, **p < 0.01, ***p < 0.001, ****p < 0.0001). **(E)** Chromatin immunoprecipitation followed by qPCR (ChIP-qPCR) analysis of LEO1, H2BK120ub, H3K9me3, and H3K27me3 occupancy at the GRP78 promoter in WT and LEO1 KO cells under glucose starvation. Statistical significance was determined by two-way ANOVA (*p < 0.05, **p < 0.01, ***p < 0.001, ****p < 0.0001). **(F)** Schematic model of GRP78 regulation in LEO1 KO cells. **(G)** Wound healing assay performed in WT and LEO1 KO cells treated with a GRP78 inhibitor under low-glucose conditions (5 mM). **(H)** qRT-PCR analysis of migration-related gene expression in HCT116 WT and LEO1 KO cells treated with a GRP78 inhibitor under glucose starvation conditions. Statistical significance was determined by two-way ANOVA (*p < 0.05, **p < 0.01, ***p < 0.001, ****p < 0.0001). **(I)** Cell viability assessed by CCK-8 assay in WT and LEO1 KO cells treated with increasing concentrations of a GRP78 inhibitor. Statistical significance was determined by two-way ANOVA (*p < 0.05, **p < 0.01, ***p < 0.001, ****p < 0.0001).

Among these factors, GRP78 emerged as a prominent stress-responsive molecule in LEO1 KO cells. GRP78 (also known as BiP) is a key ER-resident chaperone that senses misfolded proteins and regulates UPR signaling, serving as a central marker of ER stress (22). Western blot analysis demonstrated distinct ER stress response dynamics between WT and LEO1 KO cells during glucose starvation. In WT cells, GRP78 levels remained relatively unchanged, whereas pro-apoptotic ER stress markers, including ATF4 and CHOP, were markedly induced. In contrast, LEO1 KO cells showed rapid upregulation of GRP78 accompanied by delayed induction of ATF4 and CHOP (**Figure 5C**). These findings indicate that LEO1 KO cells shift ER stress responses toward adaptive rather than pro-apoptotic signaling under glucose starvation. Consistently, qRT-PCR analysis confirmed significantly greater transcriptional induction of GRP78 in LEO1 KO cells compared with WT cells during glucose starvation (**Figure 5D**).

To investigate the regulatory basis of GRP78 induction, we performed ChIP-qPCR analysis at the GRP78 promoter region. Under glucose starvation conditions, LEO1 occupancy at the GRP78 promoter increased in WT cells. In contrast, LEO1 KO cells exhibited enhanced enrichment of the active chromatin mark H2BK120ub together with reduced enrichment of the repressive histone mark H3K9me3. WT cells instead showed increased accumulation of the repressive histone modification H3K27me3 at the same promoter region. These results suggest that loss of LEO1 facilitates epigenetic activation of the GRP78 promoter under metabolic stress conditions, thereby promoting adaptive ER stress-responsive transcriptional programs (**Figure 5E**).

Given the robust induction of GRP78 in LEO1 KO cells, we next investigated its functional contribution to stress adaptation under glucose starvation. Wound healing assays demonstrated that pharmacological inhibition of GRP78 significantly reduced the migratory capacity of LEO1 KO cells under low-glucose conditions (**Figure 5G**). To further examine the transcriptional basis of this phenotype, expression of migration-associated genes was analyzed under glucose starvation conditions. GRP78 inhibition markedly attenuated induction of these genes in LEO1 KO cells (**Figure 5H**), indicating that GRP78 is required for activation of migration-related transcriptional programs during metabolic stress.

Unexpectedly, however, inhibition of GRP78 did not significantly affect cell viability in LEO1 KO cells (**Figure 5I**), despite its strong effect on migration. These findings suggest that LEO1-deficient cells maintain viability under GRP78 inhibition and preferentially utilize adaptive ER stress signaling to support migratory behavior rather than survival alone. This interpretation is consistent with the observed bias toward adaptive UPR activation and delayed induction of pro-apoptotic ER stress signaling in LEO1 KO cells.

Taken together, these results demonstrate that LEO1 deficiency promotes adaptive ER stress responses under glucose starvation conditions through epigenetic activation of GRP78 and UPR-associated pathways. Furthermore, GRP78-dependent ER stress adaptation selectively supports migration-related transcriptional programs and motility in LEO1-deficient CRC cells without substantially affecting cell viability. These findings suggest that LEO1 loss reprograms stress responses toward a migration-supportive adaptive state under metabolic stress conditions.

### Dual targeting of cholesterol metabolism and ER stress response overcomes stress adaptation of LEO1 KO cells

Given the close relationship between ER stress response adaptation and lipid metabolism (21), we next examined metabolic alterations in LEO1 KO cells. Transcriptomic analysis revealed significant upregulation of cholesterol metabolism–related pathways in LEO1 KO cells (**Figure 6A**). To investigate intracellular cholesterol dynamics, cells were treated with the lipase inhibitor orlistat, followed by analysis of BODIPY-cholesterol accumulation (40). LEO1 KO cells exhibited lower intracellular BODIPY-cholesterol signals than WT cells 12 hours after medium replacement. In contrast, orlistat treatment induced marked cholesterol accumulation in LEO1 KO cells, indicating a high-turnover cholesterol metabolic state characterized by rapid cholesterol utilization and replenishment (**Figure 6B**). Western blot analysis demonstrated reduced expression of PCSK9 in LEO1 KO cells (**Figure 6C**). Since PCSK9 negatively regulates LDL receptor–mediated cholesterol uptake (41), these findings suggest enhanced cholesterol uptake under glucose starvation. We next examined whether this metabolic rewiring generates selective vulnerability to inhibition of cholesterol biosynthesis. Treatment with atorvastatin resulted in a significantly greater reduction in viability in LEO1 KO cells compared with WT cells (**Figure 6D**), suggesting increased dependence on cholesterol metabolism to maintain stress tolerance and survival under metabolic stress conditions..

**Figure 6.**
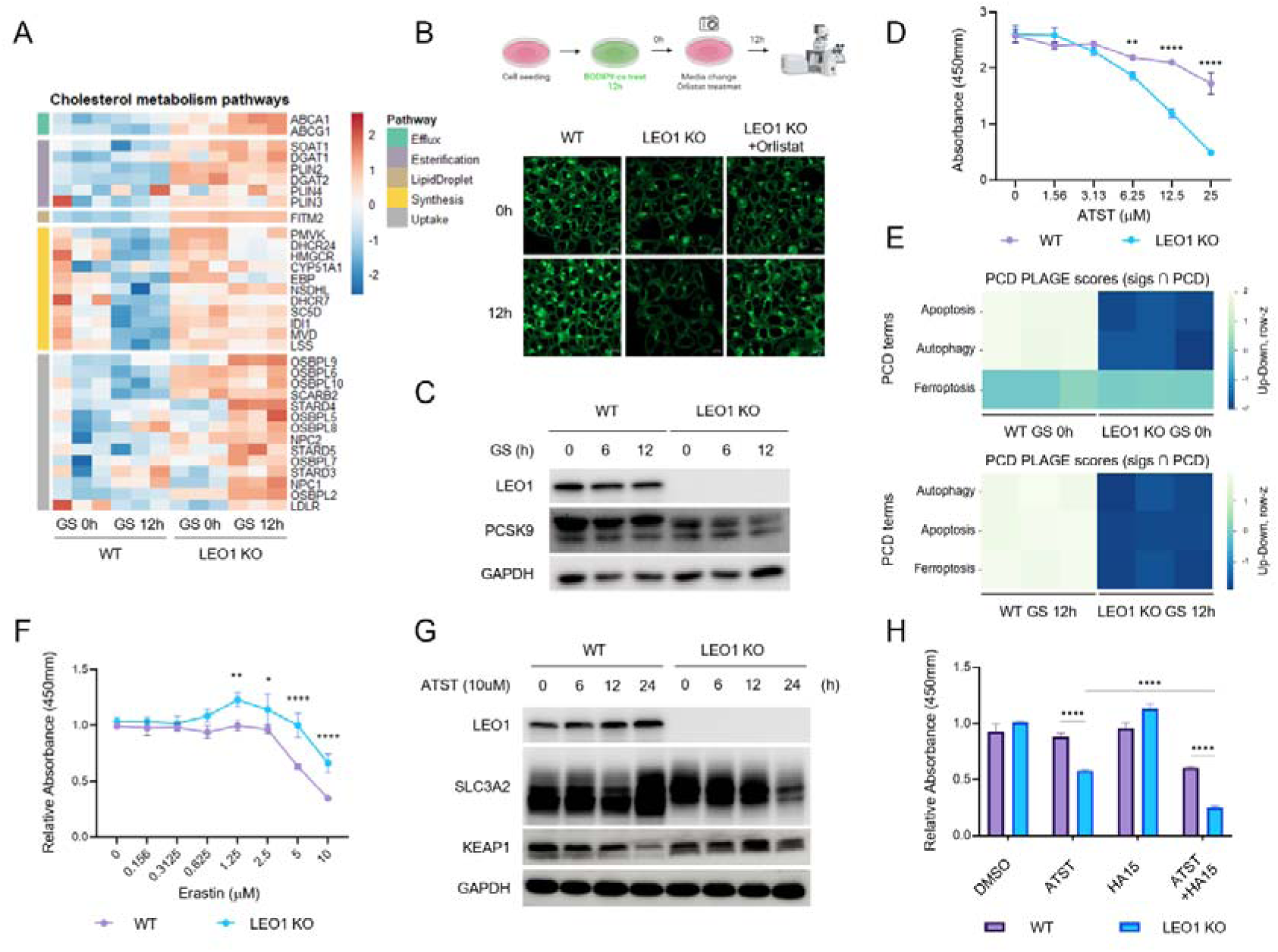
LEO1 KO cells show ferroptosis resistance and cholesterol metabolism reprogramming. **(A)** Heatmap of cholesterol metabolism-related gene expression in WT and LEO1 KO cells under glucose starvation. **(B)** Representative confocal images of BODIPY-cholesterol staining following orlistat treatment. Fluorescence signals were acquired using the 488 nm channel. Scale bars, 10 μm. **(C)** Western blot analysis of PCSK9 expression under glucose starvation conditions. **(D)** Cell viability measured by CCK-8 assay following treatment with increasing concentrations of atorvastatin. Statistical significance was determined by two-way ANOVA (*p < 0.05, **p < 0.01, ***p < 0.001, ****p < 0.0001). **(E)** PCD pathway analysis using directional PLAGE scores under glucose starvation conditions shows differential regulation of cell death programs between WT and LEO1 KO cells. **(F)** CCK-8 assay were performed to assess cell viability following treatment with increasing concentrations of ferroptosis inducer erastin. Statistical significance was determined by two-way ANOVA (*p < 0.05, **p < 0.01, ***p < 0.001, ****p < 0.0001). **(G)** Western blot analysis of ferroptosis- and redox-related proteins following atorvastatin treatment. **(H)** Cell viability was assessed following treatment with atorvastatin (ATST), GRP78 inhibitor (HA15), or their combination. Statistical significance was determined by two-way ANOVA (*p < 0.05, **p < 0.01, ***p < 0.001, ****p < 0.0001).

To further investigate stress-associated cell death pathways under glucose starvation conditions, we analyzed programmed cell death (PCD)-related transcriptional programs using the PCD database. Comparative analysis between WT and LEO1 KO cells revealed distinct regulation of ferroptosis-associated pathways, with marked alterations observed in LEO1 KO cells (**Figure 6E**). To functionally evaluate ferroptosis sensitivity, cells were treated with the ferroptosis inducers erastin. LEO1 KO cells exhibited increased resistance to erastin compared with WT cells (**Figure 6F**). These findings suggest that LEO1 loss promotes ferroptosis resistance primarily through alterations in cystine uptake–associated pathways.

To explore the associated molecular changes, expression of regulators involved in ferroptosis (42–44) and redox homeostasis (45) was evaluated. Atorvastatin treatment reduced expression of SLC3A2, a key component of the system XcL cystine transporter, whereas KEAP1 levels remained relatively unchanged (**Figure 6G**). These findings suggest that inhibition of cholesterol biosynthesis impairs cystine uptake capacity, thereby sensitizing LEO1 KO cells to stress-associated cell death.

Finally, we evaluated whether simultaneous disruption of ER stress adaptation could further enhance the therapeutic effect of cholesterol inhibition. Combined treatment with atorvastatin and the GRP78 inhibitor HA15 resulted in substantially increased cell death in LEO1 KO cells compared with either single treatment alone (**Figure 6H**). These findings indicate that co-targeting cholesterol metabolism and adaptive ER stress signaling effectively overcomes stress tolerance mechanisms in LEO1 KO colorectal cancer cells.

Taken together, these results demonstrate that LEO1 loss promotes metabolic rewiring characterized by elevated cholesterol turnover, enhanced uptake dependency, and ferroptosis resistance under metabolic stress conditions. Furthermore, LEO1 KO cells exhibit increased vulnerability to cholesterol biosynthesis inhibition, which is further potentiated by co-inhibition of GRP78-mediated ER stress adaptation. These findings identify a potential metabolic vulnerability and combinatorial therapeutic strategy in LEO1 KO colorectal cancer cells.

## Discussion

In this study, we demonstrate that loss of LEO1 reprograms transcriptional and metabolic networks to promote adaptive migration and survival under metabolic stress in CRC. Specifically, we show that LEO1 deficiency enhances ER stress adaptation through GRP78 activation, drives FOXM1-dependent migratory transcriptional programs, and induces a metabolic dependency on cholesterol metabolism. Although this cholesterol dependency may contribute to ferroptosis resistance in LEO1 KO cells, inhibition of cholesterol metabolism effectively induces cell death in these cells. These findings provide a mechanistic framework linking transcriptional regulation, stress adaptation, and metabolic rewiring in metastatic CRC (**Figure 7**).

**Figure 7.**
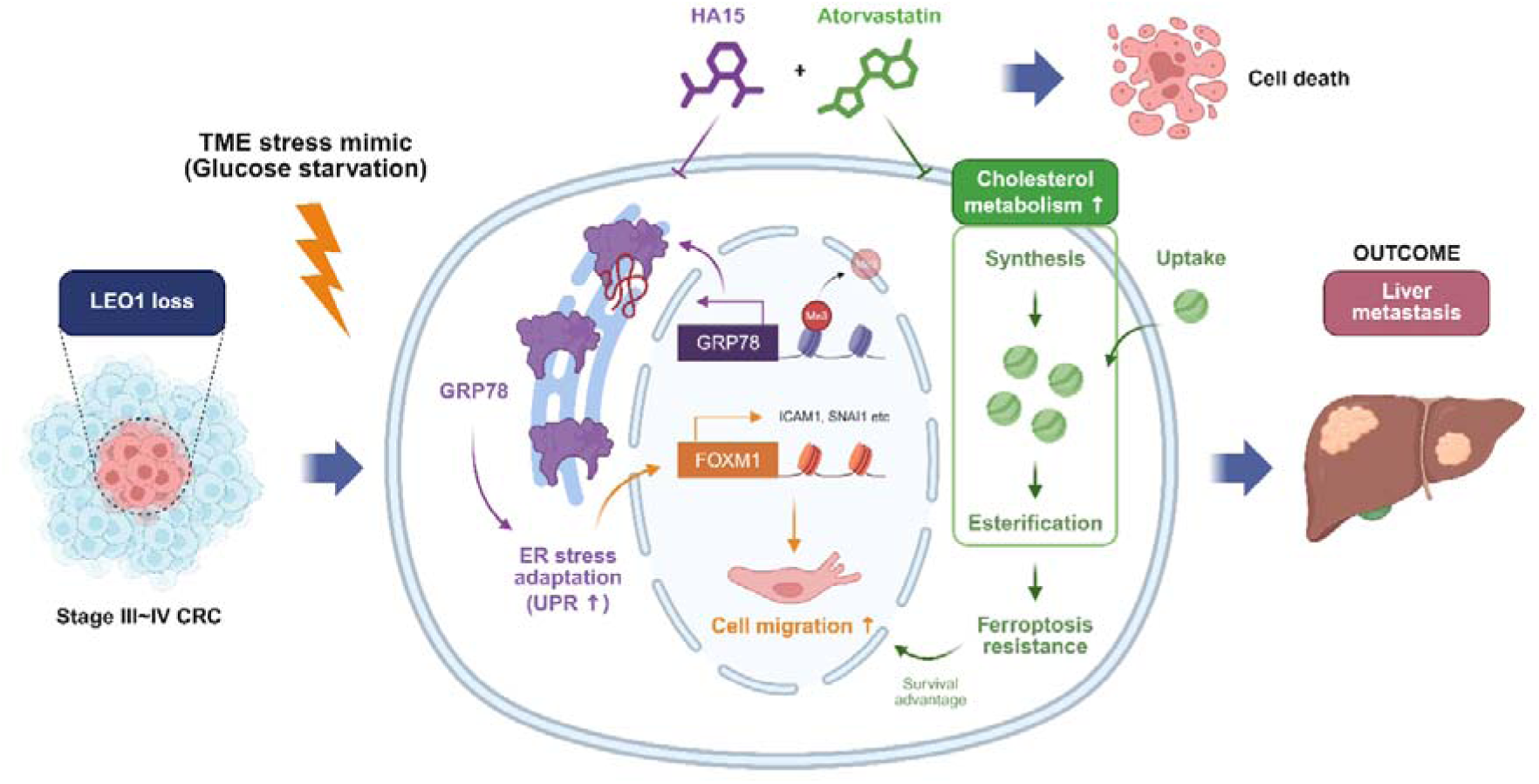
Proposed model of LEO1 loss-mediated stress adaptation and metabolic rewiring in colorectal cancer. Loss of LEO1 promotes adaptive migration and survival under metabolic stress through FOXM1-dependent migratory transcriptional programs and GRP78-mediated ER stress adaptation. In parallel, LEO1 KO cells develop increased dependency on cholesterol metabolism, which supports stress tolerance and ferroptosis resistance. Inhibition of cholesterol metabolism selectively induces cell death in LEO1 KO cells, suggesting a targetable metabolic vulnerability in metastatic CRC.

One of the key findings of this study is that LEO1 loss may confer adaptive advantages under metabolic stress, thereby contributing to clonal selection during CRC progression. While LEO1 expression is generally elevated in tumor tissues, it is paradoxically reduced in stage IV CRC and associated with poor prognosis. This suggests that loss of LEO1 may confer adaptive advantages under metastatic and metabolic stress conditions, where its loss does not simply impair cellular function but instead enables cancer cells to adopt a more aggressive phenotype. Consistent with this observation, LEO1 KO cells exhibited reduced proliferation but increased migratory capacity and nuclear enlargement, features commonly associated with metastatic potential.

Mechanistically, our data indicate that metabolic stress acts as a critical driver of LEO1-dependent phenotypic reprogramming. Under glucose starvation, LEO1 KO cells showed a marked increase in migration-associated gene expression mediated in part by the transcription factor FOXM1. FOXM1 is well known for its role in promoting proliferation and metastasis; however, our findings extend its function to stress-adaptive transcriptional regulation. Importantly, these results suggest that LEO1 normally constrains FOXM1-driven transcriptional plasticity, and that loss of LEO1 unleashes a stress-adaptive migratory transcriptional program.

In parallel, we identified GRP78-mediated ER stress adaptation as a central mechanism underlying this phenotype. Loss of LEO1 led to enhanced transcriptional activation of GRP78, particularly under glucose starvation conditions, which in turn supported both stress adaptation and migration. GRP78 is a master regulator of the UPR, and its upregulation is a well-established mechanism by which cancer cells cope with proteotoxic and metabolic stress. Our findings suggest that LEO1 loss reprograms cells to ER stress signaling while simultaneously increasing their reliance on GRP78-mediated adaptive responses. Notably, inhibition of GRP78 suppressed both migration and migration-associated gene expression programs, highlighting its functional importance in this context.

A particularly important aspect of this study is the identification of cholesterol metabolism as a metabolic vulnerability in LEO1 KO cells. Transcriptomic analyses revealed upregulation of cholesterol metabolism-related pathways, and pharmacological inhibition of cholesterol synthesis using atorvastatin selectively impaired survival of LEO1 KO cells. These findings suggest that LEO1 loss induces metabolic rewiring in which cholesterol becomes essential for maintaining membrane integrity and adaptation to stress conditions. Given the established roles of cholesterol in membrane dynamics, signaling, and oxidative stress buffering, this dependency may be particularly important for sustaining migration and survival under nutrient-deprived conditions.

Importantly, our findings highlight a functional link between ER stress adaptation, cholesterol metabolism, and ferroptosis-mediated cell death pathways, processes that are increasingly recognized as interconnected in cancer biology. ER stress responses regulate lipid synthesis and membrane homeostasis, whereas lipid metabolism influences ER function and proteostasis. In LEO1 KO cells, simultaneous activation of GRP78 and cholesterol metabolism suggests the emergence of a coordinated adaptive network that supports tumor progression. Co-targeting these pathways resulted in enhanced cell death, indicating that this dual dependency may represent a therapeutically exploitable vulnerability.

Despite these findings, several limitations should be acknowledged in this study. First, although our results demonstrate an association between LEO1 loss and metabolic dependency, the precise molecular mechanisms linking chromatin regulation to cholesterol metabolism remain to be fully elucidated. Second, while our in vitro and transcriptomic analyses provide strong support for the proposed model, *in vivo* validation using metastatic models will be necessary to confirm the physiological relevance of these findings. Finally, the clinical applicability of targeting cholesterol metabolism and ER stress pathways requires further investigation, particularly in the context of patient heterogeneity and therapeutic resistance.

In conclusion, our study identifies LEO1 as a critical regulator of stress adaptation and metabolic rewiring in CRC. Loss of LEO1 promotes GRP78-mediated migratory adaptation and cholesterol dependency, thereby creating a targetable metabolic vulnerability. These findings suggest that co-targeting ER stress adaptation and cholesterol metabolism may represent a therapeutic strategy for LEO1-low metastatic CRC.

## Materials and methods

### TCGA-COAD database

The expression levels of LEO1 between tumor and normal tissues in the TCGA Colon Adenocarcinoma (TCGA-COAD) dataset, along with stage-specific expression patterns in colorectal cancer patients, were analyzed and visualized using the GEPIA web server (http://gepia.cancer-pku.cn/), which integrates RNA sequencing data from TCGA and GTEx projects.

### Kaplan-Meier survival analysis

Survival analysis based on LEO1 expression was performed using the Kaplan-Meier Plotter database (http://kmplot.com/analysis/). Colon cancer datasets were selected, and survival was analyzed according to LEO1 expression levels using the Affymetrix probe ID 1235096_at for LEO1.

### RNA-seq Raw data processing

Raw RNA sequencing data were processed using a standard analysis pipeline. Adapter trimming and quality filtering were performed using Trimmomatic (v0.39). Read quality was assessed using FastQC (v0.11.9). Filtered reads were aligned to the human reference genome (GRCh38) using STAR (v2.7.10a). Aligned reads were sorted and processed using samtools (v1.15). Gene-level read counts were quantified using featureCounts (Subread v2.0.3) with the corresponding gene annotation file. The resulting count files were imported into R (v4.2.0), where gene-level count matrices were generated by merging individual sample files based on gene identifiers. The final count matrix was used for downstream differential expression and transcriptomic analyses.

### Differential gene expression analysis and heatmap

Gene-level count matrices were analyzed using DESeq2 in R. Lowly expressed genes were filtered prior to analysis, and differential expression analysis was performed based on the experimental design. Differentially expressed genes (DEGs) were defined using a threshold of adjusted *p*-value < 0.0001 and absolute log2 fold change ≥ 1.5. For visualization, normalized expression values were converted to log2-transformed TPM values. DEGs were mapped to gene symbols using the org.Hs.eg.db_3.20.0 annotation package, and duplicate or unmapped genes were removed. Heatmaps were generated using the pheatmap package with row scaling to visualize expression patterns across samples.

### Gene ontology (GO) enrichment analysis and GO comparison

Functional enrichment analysis was performed using the clusterProfiler package. Gene Ontology Biological Process (GO BP) terms were identified using over-representation analysis based on DEG lists. Enriched GO terms were filtered using an adjusted P value < 0.0001, and results were visualized using dot plots. Comparative GO enrichment analysis across experimental conditions was performed to identify commonly or differentially regulated biological processes. Enrichment patterns of GO BP terms were compared across datasets to identify commonly or differentially regulated biological processes.

### Gene Set Enrichment Analysis (GSEA)

GSEA was performed using the gseGO function in the clusterProfiler package. Genes were ranked based on log2 fold change values derived from DESeq2 results. Enrichment analysis was conducted for GO categories using GO Biological Process (BP) gene sets. Significantly enriched gene sets were identified using an adjusted P value < 0.0001, and results were visualized using dot plots and enrichment plots generated with enrichplot package.

### Motif enrichment analysis (cisTarget)

To identify transcription factors potentially regulating DEG-associated gene programs, motif enrichment analysis was performed using the RcisTarget package. Gene lists derived from DEGs were used as input, and enrichment of transcription factor binding motifs was assessed based on precompiled motif rankings. Candidate transcription factors were predicted based on enriched motifs associated with DEG sets.

### Clinical cohort description

Colorectal cancer tissues and matched normal tissues used for Western blot analysis were obtained from patients in the Gangnam Severance Hospital cohort. Formalin-fixed, paraffin-embedded (FFPE) tissue sections for immunohistochemical (IHC) analysis were obtained from colorectal cancer patients at Asan Medical Center. LEO1 expression was evaluated across tumor stages using IHC, including normal colon tissues (n = 3), stage I (n = 3), stage II (n = 3), stage III (n = 3) and stage IV (n = 3). In addition, matched primary stage IV tissues (n = 2) and corresponding liver metastasis tissues (n = 2) were analyzed. All human tissue samples were collected under protocols approved by the Institutional Review Boards (IRBs) of the respective institutions, and the study was conducted in accordance with relevant ethical guidelines and regulations.

### Immunohistochemistry (IHC) for patient tissues

Formalin-fixed, paraffin-embedded (FFPE) tissue sections were deparaffinized in xylene and rehydrated through a graded alcohol series to distilled water. Antigen retrieval was performed using a citrate-based antigen unmasking solution (H-3300-250, Vector Laboratories) in a pressure cooker. After heat-induced epitope retrieval, slides were cooled and washed in phosphate-buffered saline (PBS). Endogenous peroxidase activity was quenched using BLOXALL blocking solution (SP-6000-100, Vector Laboratories) for 10 min, followed by washing in PBS. Sections were then incubated with normal blocking serum for 20 min to reduce nonspecific binding. Primary antibodies were applied and incubated at room temperature for 30 min. For LEO1 staining, an anti-LEO1 antibody for IHC (Antibodies, A46943) was used at a 1:200 dilution. After washing, sections were incubated with biotinylated secondary antibody (BA-1000-1.5, Vector Laboratories) for 30 min, followed by incubation with the VECTASTAIN Elite ABC-HRP reagent (PK-6102, Vector Laboratories) for 30 min according to the manufacturer’s instructions. For signal detection, sections were incubated with 3,3′-diaminobenzidine (DAB) substrate (SK-4100, Vector Laboratories) until appropriate staining intensity developed. The reaction was stopped by rinsing in water, and sections were counterstained with hematoxylin, dehydrated, cleared, and mounted. Deparaffinization, rehydration, and dehydration of tissue sections were carried out by the Histology Core Facility at the KAIST Graduate School of Medical Science.

### Hematoxylin and Eosin staining

Tissue sections were deparaffinized in xylene (3 changes, 2 min each) and rehydrated through a graded ethanol series (100% ethanol, 3 changes, 2 min each; followed by 95%, 90%, 80%, and 70% ethanol, 2 min each) and then rinsed in distilled water for 2 min. Sections were stained with hematoxylin for 5 min and washed in distilled water for 2 min. Differentiation was carried out using 1% acid alcohol (1% HCl in 70% ethanol) for 5 seconds, followed by washing in distilled water for 2 min. Bluing was performed in 0.4% ammonia water for 2 min and rinsed in distilled water for 2 min. Sections were then counterstained with eosin for 1 minute. After staining, sections were dehydrated through an ascending ethanol series (70%, 80%, 90%, 95%, and 100% ethanol; 2 min each), with 100% ethanol applied in three changes (2 min each), followed by clearing in xylene (4 changes, 2 min each) and mounted for microscopic analysis.

### Cell culture

Human colorectal cancer cell lines were cultured in Roswell Park Memorial Institute medium (RPMI-1640, Welgene, LM011-01) supplemented with 10% fetal bovine serum and 1% penicillin/streptomycin (PS, Welgene, LS203-01). The normal human colon epithelial cell line HCEC-1CT was cultured in Colon ColoUp ready-to-use medium (EVERCYTE, MHT-039) according to the manufacturer’s instructions. All cells were maintained at 37°C in a humidified incubator with 5% COL. For glucose starvation experiments, cells were cultured under standard conditions, washed with phosphate-buffered saline (PBS), and subsequently incubated in RPMI-1640 medium without glucose (Welgene, LM011-60) supplemented with 10% dialyzed FBS (Gibco, 26400-044) and 1% PS. Cells were seeded in T25 flasks and, upon reaching the desired confluency, the culture medium was replaced with starvation medium. Hypoxic conditions were maintained at 1% OL using a HERAcell 150i incubator (Thermo Fisher Scientific).

### Construction of sgRNA expression vector

sgRNA oligonucleotides targeting LEO1 (Fwd: 5′-CACCGCGGATATGGAGGATCT CTT-3′; Rev: 5′-GTAGAAGAGATCCTCCATATCCGC-3′) were synthesized and resuspended in annealing buffer (10 mM Tris-HCl, pH 7.5-8.0, 50 mM NaCl, 1 mM EDTA). The oligos were mixed at equimolar concentrations (2 μg each in a total volume of 50 μl), incubated at 95°C for 2 min, and gradually cooled to room temperature over 45 min for annealing. The annealed oligos were diluted (5 μl in 45 μl nuclease-free water) prior to ligation. The lenti-sgRNA puro vector (Addgene plasmid #104990) was digested with BsmBI-v2 (NEB) and purified, followed by ligation of the annealed oligos using T4 DNA ligase (Enzynomics, M001L) according to the manufacturer’s instructions.

### Construction of shRNA expression vector

For FOXM1 KD, shRNA oligonucleotides targeting FOXM1 (Fwd: 5′-CCGGGCCAATCGTT CTCTGACAGAACTCGAGTTCTGTCAGAGAACGATTGGCTTTTTG-3′; Rev: 5′-AATTC AAAAAGCCAATCGTTCTCTGACAGAACTCGAGTTCTGTCAGAGAACGATTGGC-3′) were synthesized and resuspended in annealing buffer. The oligos were mixed at equimolar concentrations and annealed by incubation at 95°C for 2 min followed by gradual cooling to room temperature over 45 min. The annealed oligos were diluted prior to ligation. The pLKO.1-puro vector (Addgene plasmid #8453) was digested with AgeI and EcoRI, purified, and ligated with the annealed oligos using T4 DNA ligase according to the manufacturer’s instructions.

### Lentiviral transduction for generation of knockout and knockdown cells

Lentiviral particles were produced using Lenti-X™ 293T cells, which were seeded in 100 mm culture dishes (Sarstedt, 83.3902). For transfection, 3 μg of psPAX2 (Addgene plasmid #12260), 1 μg of pMD2.G (Addgene plasmid #12259), and 4 μg of lentiviral transfer vector were mixed with 20 μl of 1 mg/ml polyethyleneimine (PEI; Merck, 408727) and 472 μl of DMEM. The mixture was incubated at room temperature for 15 min and then added to the cells for transfection. At 24 h post-transfection, the medium was replaced with 10 ml of fresh RPMI- 1640, and the cells were further incubated at 37°C in a 5% COL incubator for an additional 24 h. The culture supernatant containing lentiviral particles was then collected and filtered through a 0.45 μm filter (GVS, FJ25ASCCA004FL01). For transduction, the filtered viral supernatant was supplemented with polybrene (10 μg/ml; Merck, TR-1003-G) and added to target cells.

### Western blot analysis

Proteins from cells and tissues were detected by Western blot analysis. Cells and tissues were washed with 1× phosphate-buffered saline (PBS, Welgene, LB001-02) and lysed in EBC200 buffer (50 mM Tris-HCl, pH 8.0, 200 mM NaCl, 0.5% NP-40) supplemented with protease inhibitor cocktail (Enzynomics, P3100-05). Samples were sonicated once at 30% amplitude for 20 sec and centrifuged at 13,000 rpm for 10 min at 4°C. The supernatants were transferred to new tubes, and protein concentrations were quantified using the Bradford assay. Equal amounts of protein (20 μg) were denatured in SDS sample buffer at 100°C for 10 min. Protein samples were separated on polyacrylamide gels and transferred to PVDF membranes (Millipore, IPVH00010). Membranes were blocked with 5% skim milk for 45 min at room temperature and incubated with primary antibodies overnight at 4°C. Membranes were washed with 1× TBST and incubated with HRP-conjugated secondary antibodies for 1 h at room temperature. Protein bands were visualized using enhanced chemiluminescence (ECL) and detected using the ImageQuant LAS500 system (GE Healthcare). The following primary antibodies were used: anti-LEO1 (Invitrogen, A300-175A), anti-GAPDH (Cell Signaling Technology, 2118S), anti-LAMIN A/C (Cell Signaling Technology, 2032S), Cell Cycle Regulation Antibody Sampler Kit (Cell Signaling Technology, 9932T), anti-BiP (Cell Signaling Technology, 3177T), anti-CHOP (Cell Signaling Technology, 2895T), anti-ATF4 (Cell Signaling Technology, 11815T), and anti-PCSK9 (Cell Signaling Technology, 85813S).

### RNA precipitation

Total RNA was extracted using TRIzol reagent (Invitrogen, 15596026) according to the manufacturer’s instructions. Cells were washed with PBS and lysed in 1 ml TRIzol reagent. Phase separation was performed by adding 200 μl chloroform followed by centrifugation at 12,000 × g for 10 min at 4°C. The aqueous phase was transferred to a new tube, mixed with 500 μl isopropanol, and incubated for 10 min at room temperature. Samples were centrifuged at 12,000 × g for 10 min at 4°C, and RNA pellets were collected. Pellets were washed with 1 ml of 75% ethanol, air-dried for approximately 10 min, and dissolved in RNase-free water. Dissolved RNA samples were incubated at 65°C for 10 min before use.

### Real time PCR (qRT-PCR)

To analyze mRNA expression levels, cDNA was synthesized from total RNA using a reverse transcription kit (Enzynomics, EZ005S). Quantitative real-time PCR (qRT-PCR) was performed using SYBR Green master mix (Enzynomics, RT500M) on a QuantStudio 1 system (Applied Biosystems). Relative mRNA expression levels were calculated using the ΔΔCt method. All experiments were performed in triplicate, and statistical analyses were conducted using Prism 9 software. The primer sequences used for quantitative real-time PCR were as follows: LEO1 (Fwd: 5′-AGAAGCGGATAGTGACACTGAGGT-3′; Rev: 5′-TTCATCAACA GGCTGTCCTGGAGT-3′), SNAI1 (Fwd: 5′-TCGGAAGCCTAACTACAGCGA-3′; Rev: 5′-AGATGAGCATTGGCAGCGAG-3′), ICAM1 (Fwd: 5′-AGCGGCTGACGTGTGCAGTAAT -3′; Rev: 5′-TCTGAGACCTCTGGCTTCGTCA-3′), MYL9 (Fwd: 5′-GGATGTGATTCGCA ACGCCTTTG-3′; Rev: 5′-CGGTACATCTCGTCCA CTTCCT-3′), LAMA1 (Fwd: 5′-GAA GGTGACTGGCTCAGCAAGT-3′; Rev: 5′-AGGCGTC ACAACGGAAATCGTG-3′), HSPA5 (Fwd: 5′-GCCGTCCTATGTCGCCTTC-3′; Rev: 5′-TGGCGTCAAAGACCGTGTTC -3′), and FOXM1 (Fwd: 5′-TCTGCCAATGGCAAGGTCTCCT-3′; Rev: 5′-CTGGATTCGGT CGTTTCTGCTG-3′). The primer sequences used for ChIP-qPCR were as follows: HSPA5 (Fwd: 5′-GATAACATCCGCCCCATCCG-3′; Rev: 5′-GAGTG AAG GCGGGACTTGTG-3′).

### CCK-8 assay

To assess the cell proliferation and viability, cells were seeded at 1,000-2,000cells/well for proliferation and 5,000cells/well for viability. Following cell seeding and drug treatment, CCK-8 solution (Dojindo, Cell Counting Kit-8) diluted in culture medium at a 1:9 ratio was added to each well. Plates were incubated for 1h 30min-2h in incubator. Absorbance at 450 nm was measured by BioTek Synergy H1 Microplate Reader. All experiments were performed in triplicate, and statistical analyses were conducted using Prism 9 software.

### Transwell assay

Cell migration was assessed using 6.5 mm Transwell® inserts with 8.0 μm pore size in 24-well plates (Corning, 3422). HCT116 wild-type (WT) and LEO1 knockout (KO) cells were trypsinized and resuspended in serum-free medium. A total of 5 × 10L cells in 100 μl were seeded into the upper chamber. The lower chamber was filled with RPMI-1640 medium supplemented with 10% fetal bovine serum (FBS) as a chemoattractant. Cells were incubated at 37°C in a humidified atmosphere with 5% COL for 24 h. After incubation, non-migrated cells remaining on the upper surface of the membrane were gently removed using a cotton swab. Migrated cells on the lower surface were fixed with 100% methanol for 30 min and subsequently stained with 0.25% crystal violet for 30 min. Excess stain was removed by washing with tap water. Stained cells were visualized using an EVOS XL core.

### Wound scratch assay

To assess cell migration, cells were seeded at 5 × 10L-7 × 10L cells/well in ImageLock 96-well plates (Sartorius). After 24 h incubation, wound scratches were generated using the IncuCyte WoundMaker tool according to the manufacturer’s instructions. Culture medium was replaced with medium corresponding to each experimental condition. Wound closure was monitored using the IncuCyte live-cell imaging system, and relative wound density was quantified using the IncuCyte Scratch Wound Analysis Software Module.

### Immunofluorescence staining

Cells were seeded in 35 mm confocal dishes at a density of approximately 3 × 10L cells. Upon reaching the desired confluency, cells were fixed with 4% paraformaldehyde for 15 min at room temperature, followed by washing with PBS twice for 10 min each. Cells were then permeabilized with 0.1% Triton X-100 in PBS (PBST) for 10 min at room temperature and washed twice with PBST. For blocking, cells were incubated in blocking buffer containing 5% bovine serum albumin (BSA) and 5% normal donkey serum for 1 hour at room temperature. Primary antibodies were diluted in blocking buffer according to the manufacturer’s instructions and incubated with the cells overnight at 4°C. After washing three times with PBST, cells were incubated with fluorescence-conjugated secondary antibodies diluted in blocking buffer for 1 hour at room temperature in the dark. Following three washes with PBST, nuclei were stained with Hoechst 33342 (1:5000 in PBS). Cells were washed three additional times with PBS and imaged using a confocal laser scanning microscope (LSM980, Carl Zeiss).

### Flow cytometry for cell cycle analysis

Cells were trypsinized using TE buffer and counted to obtain a concentration of 3 × 10L cells/ml in a total volume of 4 ml. Cells were centrifuged at 1,000 rpm for 10 min, and the pellets were washed with PBS followed by additional centrifugation. For fixation, cell pellets were vortexed while 70% ethanol was slowly added dropwise to ensure proper mixing. Fixed cells were mixed with 3% BSA/PBS at a 1:1 ratio, inverted gently, and centrifuged at 800 rpm for 5 min. The supernatant was removed, and pellets were washed again with 3% BSA/PBS. Cells were centrifuged at 800 rpm for 5 min and resuspended in nucleic acid staining solution containing RNase A (100 μg/ml in PBS). Samples were incubated at room temperature for 20 min protected from light. Cells were filtered through a Multi C-strainer (SPL94040) into FACS tubes prior to staining. For propidium iodide (PI) staining, 10 μl PI solution (1 mg/ml) was added and mixed thoroughly. Cell cycle distribution was analyzed using an LSRFortessa X-20 flow cytometer.

### Geminin based cell cycle analysis via lentiviral transduction

For geminin-based cell cycle analysis, HCT116 WT and LEO1 KO cells were transduced with a CSII-EF-mCherry-hGeminin lentiviral vector. Lentiviral particles were produced in HEK293T cells by co-transfection of psPAX2, pMD2.G, and CSII-EF-mCherry-hGeminin plasmids using polyethyleneimine (PEI). After 48 h, viral supernatants were collected, filtered, and transferred to target cells in the presence of polybrene. Following infection, cells were trypsinized, serially diluted, and seeded into 96-well plates for clonal expansion over 1–2 weeks. Single-cell clones expressing mCherry-Geminin were monitored using the dilution cloning function of the IncuCyte live-cell imaging system. Cell cycle progression was measured and quantified using the IncuCyte system.

### Nuclear area measurement

Cells were seeded under the same conditions as described for immunofluorescence experiments. Nuclear staining was performed using Hoechst 33342. Images were acquired, and nuclear area was quantified using ZEN software (version 3.6, ZEN Lite, Zeiss). Data were analyzed using GraphPad Prism 9 software. Statistical significance was determined using an unpaired t-test.

### Chromatin immunoprecipitation (ChIP)

Magnetic beads were washed twice with dilution buffer, and antibodies were preincubated at room temperature for 3 h. Cells were collected by treating them with trypsin, followed by centrifugation at 1000 rpm for 3 min, and then resuspended in 6 ml of media. The cells were fixed with 1% formaldehyde at room temperature for 10 min and quenched with 125 mM glycine at room temperature for 5 min. After centrifugation at 750 rcf at 4°C, the supernatant was removed, and the cells were washed twice with 6 ml of cold PBS. The supernatant was removed, and 3×10^6^ cells per condition were counted. The cells were divided and resuspended in 300 μl of high salt lysis buffer (0.8 M NaCl, 25 mM Tris pH7.5, 5 mM EDTA pH8.0, 1% Triton X-100, 0.1% SDS, 0.5% sodium deoxycholate, 1X protease inhibitor cocktail), followed by sonication with 30% amplification and 30-sec pulses for 10 cycles. The sample was then centrifuged at 13,600 rcf for 30 min at 4°C, and the supernatant was transferred to a new tube. Dilution buffer (25 mM Tris pH7.5, 5 mM EDTA pH8.0, 1% Triton X-100, 0.1% SDS, 1X protease inhibitor cocktail) was added to 300 μl of the sample about 1 ml, and 65 μl of input was kept in a new tube. Magnetic beads were added to the sample about 20 μl, and the mixture was incubated overnight on a rotator at 4°C. The sample was washed once each with Buffer A (140 mM NaCl, 50 mM HEPES pH7.9, 1 mM EDTA pH8.0, 0.1% SDS, 0.1% sodium deoxycholate), Buffer B (500 mM NaCl, 50 mM HEPES pH7.9, 1 mM EDTA pH8.0, 1% Triton X-100, 0.1% SDS, 0.1% sodium deoxycholate), and Buffer C (20 mM Tris pH7.5, 1mM EDTA pH8.0, 250 mM LiCl, 0.5% NP-40 alternative, 0.5% sodium deoxycholate), followed by washes twice with TE buffer (10 mM Tris pH7.5, 1 mM EDTA pH8.0), each for 5 min at room temperature. 100 μl of elution buffer (10 mM Tris pH7.5, 1 mM EDTA pH8.0, 1% SDS) was added to the beads, and the mixture was incubated at 65°C for 5 min and rotated at room temperature for 15 min, repeating this process twice. The eluted sample was transferred to a new tube, and 135 μl of elution buffer was added to the input. To each sample, 8 μl of 4 M NaCl and 0.4 μl of 10 mg/ml RNase A were added, and the mixture was incubated at 65°C overnight to reverse the protein-DNA crosslinking. Subsequently, 0.2 μl of 0.5 M EDTA and 2 μl of 20 mg/ml proteinase K were added, and the mixture was incubated at 45°C for 2 h. DNA was purified using the Qiagen DNA purification kit, and the results were analyzed by qRT-PCR.

### BODIPY-cholesterol uptake and orlistat treatment

Cells were seeded onto confocal imaging plates (SPL, 200350) and allowed to attach under standard culture conditions. To assess cholesterol utilization, cells were incubated with BODIPY-cholesterol (MCE, HY-125746) (10 μM) for 12 hours. After incubation, cells were washed with PBS to remove excess probe and then replaced with either fresh complete medium or complete medium supplemented with orlistat (Merck, O4139) (20 μM). After 12 hours of incubation, fluorescence images were acquired using a LSM980 confocal microscope (Zeiss) with the GFP channel.

### Programmed cell death-related gene expression analysis

Differential gene expression analysis was performed between WT_0h and KO_0h samples using DESeq2. Genes with an adjusted P value < 0.05 and an absolute log2 fold change ≥ 1.5 were considered significantly differentially expressed. To evaluate programmed cell death-related transcriptional changes, programmed cell death gene annotations were obtained from RCDdb. Genes annotated in RCDdb were categorized according to programmed cell death subtypes, and genes associated with autophagy-dependent cell death, apoptosis, and ferroptosis were selected for downstream analysis. Significant differentially expressed genes from the WT_0h versus KO_0h comparison were then intersected with the selected RCDdb-derived programmed cell death gene sets. For visualization, normalized expression values of the overlapping programmed cell death-related genes were extracted across WT_0h, WT_12h, KO_0h, and KO_12h samples. Expression values were Z-score normalized across samples for each gene to emphasize relative expression differences among conditions. Heatmaps were generated using hierarchical clustering of genes, while the sample order was manually fixed according to the experimental design. Genes were further annotated based on their programmed cell death category and direction of differential expression in KO_0h relative to WT_0h.

